# Targeting *Plasmodium falciparum* calcium dependent protein kinase-1: Computational modelling approach towards the discovery of antimalarial drug from medicinal herbs and clinically approved anti-malarial drugs pharmacophores

**DOI:** 10.1101/2022.08.22.504824

**Authors:** Temitope Isaac Adelusi, Taiwo Ooreoluwa Ojo, Bolaji Olawale Quadri, Precious Moyosoluwa Oyewole, Elijah Kolawole Oladipo

## Abstract

The Key participants in the calcium signaling process in the malaria parasite are calcium dependent protein kinases. It has been established that *Plasmodium falciparum* calcium dependent protein kinase 1 (*Pf*CDPK1) is essential for the invasion of host erythrocytes as well as for the parasite’s asexual development. Due to malaria’s resistance to currently available anti-malarial medications, it has become necessary to create effective inhibitory agents that target invasion-dependent proteins. In this study, libraries of 856 chemicals were screened using a variety of computational methods. Physicochemical property testing, molecular docking analysis, and pharmacokinetic ADMET profiling were used to determine these drugs’ capacity to inhibit *Pf*CDPK1. 88 of the 856 compounds derived from known antimalarial botanicals and clinically approved antimalarial drug pharmacophores have demonstrated a potent inhibitory effect on *Pf*CDPK1. Additionally, based on research done on the purported inhibitory standard of *Pf*CDPK1 (Purfalcamine), which had a binding affinity of −8.8 KJ/mol, twelve compounds, which include 17-hydroxyazadiradione, Nimbaflavone, Dehydrocarpaine-II, Picracin, Epicatechin gallate, and Naringin, ZINC000012504271, ZINCC000100029905, ZINC000004149268, ZINC000078832372, ZINC000102739374 and ZINC000198562314 had exhibited better binding affinities (−8.8, −8.8, −9.2, −9.1, −9.3, −10.0, −8.1, −7.9, −7.9, −8.2, −8.3, −7.9 KJ/mol respectively), pharmacokinectics, and physicochemical properties. According to our findings, 88 compounds are likely pharmacological targets with domiciled *Pf*CDPK1 inhibitory effects. Nevertheless, our findings highlight these twelve stand-out compounds, suggesting that they may be the best candidates with regard to therapeutic efficacy.

## INTRODUCTION

*Plasmodium falciparum*, which causes malaria, is a parasitic disease spread by mosquitoes that affects 200-400 million people and kills 400,000 people every year (mostly children in sub-Saharan Africa) (Bansal et al., 2013; Wang et al., 2020). Malaria continues to be a serious public health issue in tropical regions despite ongoing efforts to control it. In recent years, the emergence and spread of artemisinin-resistant parasites has caused grave concerns about the availability of efficient malaria prevention methods (Iyer et al., 2018). Consequently, there is an urgent need for novel antimalarial medication lines (Lemercier et al., 2009). The complex life cycle of the human malaria parasite *Plasmodium falciparum* includes infection of both the human host and the vector. It begins to spread in the liver before invading the erythrocytes, where it then develops. Recent research has shown that practically all stages of parasite growth involve intracellular signaling (Iyer et al., 2018)

Erythrocyte invasion is a multistep, intricate process that is made easier by several interactions between host receptors and parasite ligands. The micronemes and rhoptries, two apical organelles of Plasmodium merozoites, are home to several parasite proteins implicated in these interactions. Since invasion depends on the timely release of these proteins to the merozoite surface to enable receptor engagement, a thorough understanding of the molecular mechanisms underlying the malaria parasite’s signaling pathways may shed light on previously unknown aspects of the parasite’s biology and facilitate the targeting of important enzymes for drug development (Ahmed et al., n.d.; Bansal et al., 2013).

The family of gene products known as calcium-dependent protein kinases is found in plants and alveolates but not in humans. The genome of Plasmodium falciparum contains five genes that code for the five CDPKs (*Pf*CDPK1–5). Amino acid identity varies between 39 and 56% among paralogues of the CDPK family (Bansal et al., 2017; Green et al., 2008; Lemercier et al., 2009). *Pf*CDPK1 was initially discovered by Zhao et al. It has been suggested that the protein is connected to the parasitophorous vacuole membrane (PVM). Expression of *Pf*CDPK1 is only present in late blood stages of *Plasmodium falciparum* throughout the asexual cycle, and it is highest in mature schizonts (Green et al., 2008).

An *in vitro* phosphorylation test with *Pf*CDPK1 demonstrated that it phosphorylates the motor complex’s constituent parts (Bansal et al., 2017). About 30 hours after invasion, *Pf*CDPK1 is expressed throughout the parasite’s intraerythrocytic stage and is linked to membrane and organelle fractions (Holder et al., 2012). *Pf*CDPK1 buildup within the parasite’s cells in particular compartments raises the possibility that it plays a role in the active membrane biogenesis of blood stage parasites as well as the mechanism of merozoite invasion (Lemercier et al., 2009). The fact that *Pf*CDPK1 is present throughout the growing schizont and in the free merozoite suggests that it may perform a variety of functions both within the growing intracellular parasite and during the invasion of host erythrocytes by merozoites. These functions may be regulated by calcium flux waves within the cell, particularly within the pellicle, as well as by post translational modification (Holder et al., 2012).

During erythrocyte merozoite invasion and intracellular development, CDPK1 may be differentially phosphorylated by autophosphorylation, the action of other kinases and/or phosphatases, and other processes. This could affect the enzyme’s activity and substrate selectivity (Holder et al., 2012). Its involvement in host invasion may be explained by the phosphorylation of *Pf*GAP45 and *Pf*MTIP, two glideosome assembly components (Iyer et al., 2018). Because parasites treated with CDPK inhibitors exhibit less gliding and invasion in plasmodium, it is possible that the substrates of CDPKs are directly engaged in motility and erythrocyte invasion (Green et al., 2008). Merozoites could not exit mature segmented schizonts when purfalcamine, a known inhibitor of *Pf*CDPK1, was use as a treatment. Later research showed that purfalcamine treatment of parasites prevented microneme release and subsequent invasion of host RBCs (Bansal et al., 2013, 2017; Green et al., 2008).

Through our investigation of *Pf*CDPK1 and its function in host cell invasion, we may have been able to identify more effective inhibitors of *Pf*CDPK1 with favorable ADMET profile scores, greater binding affinities, and favorable drugability parameters. This study used a variety of computational and bioinformatics methodologies to carry out virtual screening. High-throughput screening (HTS), which looks for hits in the early stages of drug research, is complemented by virtual screening. Specifically, compounds are required to launch efforts toward a clinical candidate after a macromolecular target is chosen. *Pf*CDPK1 was used as the target macromolecule in the investigation of a library of compounds from plants that have been reported to have anti-malarial properties and pharmacophores of available anti-malaria drug active compounds. Our aim is to propose compounds with good pharmacokinetic, pharmacodynamic and physicochemical profile coupled with inhibitory activity towards *Pf*CDPK1 target, which may be channeled into preclinical and clinical trials. These compounds might later emerge as drugs/nutraceuticals for the management/treatment of malaria.

## MATERIALS AND METHOD

### Preparation of target protein and validation

The National Center for Biotechnology Information (NCBI), which may be accessed at (https://www.ncbi.nlm.nih.gov/), provided the *Pf*CDPK1 protein amino acid sequence in FASTA format with the accession number P62344 (Sanami et al., 2021). To create the 3D structure of the protein, the protein’s 524 amino acid residues were uploaded to the I-TASSER server (https://zhanggroup.org/I-TASSER/) (Roy et al., 2010). The I-TASSER site uses a cutting-edge algorithm to create a 3D structure that most correctly depicts the protein’s structure and its functions (Samad & Ahammad, 2020). The top five models of the provided protein sequence, as well as the C-score, TM-score value, RMSD, and other metrics, can be predicted and determined by this online service. In accordance with the C-score value, the generated 3D structure was downloaded in the PDB format. The server has a C-score from −5 to 2, where a higher number denotes a highly confident protein model (Adhikari et al., 2018). The validation of the modeled protein structure is crucial. The projected tertiary structure was validated using ProSA-web (https://prosa.services.came.sbg.ac.at/prosa.php) to identify any potential inaccuracies (Wiederstein & Sippl, 2007). The server offers details on the protein structure’s overall quality score and any problematic regions (Onile et al., 2020). By using the Ramachandran plot server (https://zlab.umassmed.edu/bu/rama/), the improved tertiary structure was verified(Anderson et al., 2005).The structure of the protein was examined, and a Ramachandran plot was created. The protein structure was then submitted to Autodock Vina v1.5.7, where the PDBQT file was created by editing the Histidine molecules and adding Kollman charges and hydrogen bonds (Trott & Olson, 2011).

### Determination of target protein active site

Knowing the amino acid residues of a protein’s active site is crucial for identifying the site at which ligands will interact with the target protein (Prada-gracia et al., 2016). Utilizing the CASTp webpage (http://sts.bioe.uic.edu/castp/index.html?201l), it was possible to detect the amino acid residues that would be found in the protein’s active pocket (Tian et al., 2018).

### Ligand selection and preparation

Ethnopharmacological study was done on reported plants with potential ameliorative effect on *Plasmodium falciparum*. Nine (9) plants were selected namely; *Anacardium ocidentale* (Ross., n.d.; Odoh et al., 2018; Oladeji et al., 2020; Olorunnisola et al., 2013; Omosun et al., 2013; Salehi et al., 2019), *Carica papaya* (Al-seadi et al., 2021; Dogara & Labaran, 2020; Odoh et al., 2018; Oladeji et al., 2020; Olorunnisola et al., 2013; Omosun et al., 2013), *Citrus aurantofolia* (Odoh et al., 2018; Oladeji et al., 2020; Omosun et al., 2013), *Morinda lucida* (Idowu et al., 2010; Odoh et al., 2018; Okoh et al., 2011; Oladeji et al., 2020; Olorunnisola et al., 2013), *Azadirachta indica* (Alzohairy, 2016; Dogara & Labaran, 2020; Idowu et al., 2010; Odoh et al., 2018; Oladeji et al., 2020; Olorunnisola et al., 2013; Omosun et al., 2013; Ugboko et al., 2020), *Alium sativum* (Lifongo et al., 2014; Odoh et al., 2018; Oyeyemi et al., 2019), *Pikralima nitida* (Campos et al., 2020; Aghedo et al., 2021; Erharuyi et al., 2014; Lifongo et al., 2014; Oyeyemi et al., 2019), *Mangnifera indica* (Dogara & Labaran, 2020; Idowu et al., 2010; Maldonado-celis, 2019; Sonia et al., 2010; Odoh et al., 2018; Oladeji et al., 2020; Olorunnisola et al., 2013; Omosun et al., 2013; Kumar et al., 2021), *Chromalaena odorata*(Idowu et al., 2010; Odoh et al., 2018; Oladeji et al., 2020; Olorunnisola et al., 2013; Vijayaraghavan et al., 2017; Yusuf et al., 2021) and their respective phytochemicals were retrieved from PUBCHEM (https://pubchem.ncbi.nlm.nih.gov/) in SDF format (Kim et al., 2016)and were later converted to PDB file using Biovia discovery studio. To prepare for docking, the PDBQT file was prepared when the PDB file was submitted to Autodock Vina v1.5.7 (Trott & Olson, 2011). Artemether, Quinine, Chloroquine, Atovaquone, Primaquine, Mefloquine, Tafenoquine, and Doxycycline were among the active ingredients of malaria medications that were subjected to pharmacophoric modeling. Their pharmacophores were generated using ZINC15 (https://zinc15.docking.org/) in SDF format(Sterling & Irwin, 2015), and they were prepared similarly to the previous batch. Purfalcamine, a reported PFCDPK1 inhibitor, was also obtained in SDF format from pubchem and prepared for analysis.

### ADMET and drug-likeness profiling

Molinspiration Online tools (https://molinspiration.com/) was used to forecast parameters including molecular weight, milogP, number of hydrogen hydrogen bond donors, number of hydrogen bond acceptors, and number of violations in order to evaluate the drug-likeness of the chosen phytochemical while ADMETlab 2.0 server (https://admetmesh.scbdd.com/) was used to assess the phytochemical’s pharmacokinetics features, including absorption, distribution, metabolism, excretion, and toxicity in order to help choose the optimal therapeutic candidate (Xiong et al., 2021).

### Molecular Docking Procedure

Autodock v1.4.7 was used for additional analysis in identifying the specific interactions between the chosen phytochemicals and the target receptor. Active sites were identified, and a grid box with the dimensions 59×89×89, binding center of 63.458, 59.662, and 83.396 x, y, and z respectively were created. Docking was carried out using a Microsoft Windows 10 pro computer. Each target’s binding affinity as well as the bonds involved in the interaction were assessed and reported.

## RESULT

### Preparation of target protein and validation

I-TASSER server’s 3D homology modeling prediction offered the top 5 models. Out of the five models, the one with the greatest C-Score (−1.03) was picked for further analysis. Figure 2 shows a structural depiction of the predicted structure. The Ramachandran plot server and ProSA web server were used to validate the 3D structure after modeling. The Ramachandran plot server’s results showed that 87.93% of the amino acids are present in the highly preferred observation region, 7.0% in the preferred observation region, and only 4.98% in the questionable observation region. A z-score value of −7.17 was predicted by the ProSA web server for the tertiary structure. Figure 3 illustrates the results of the tertiary structure validation.

**Figure 2:**
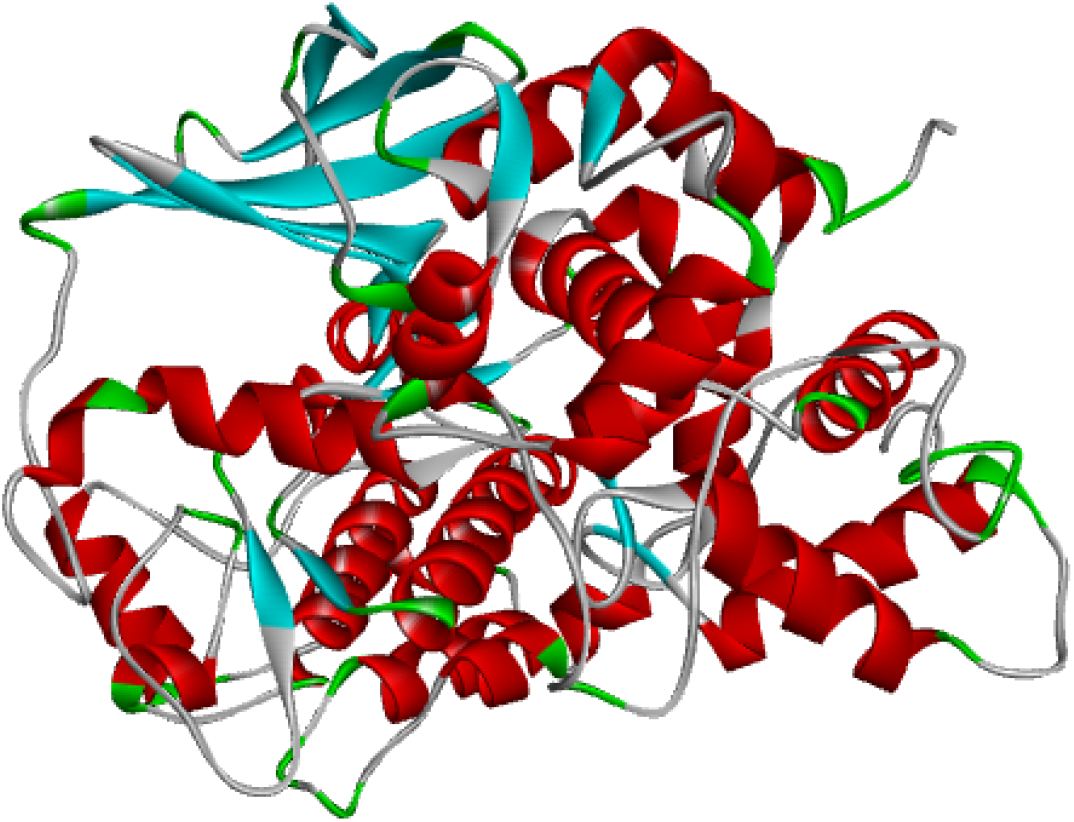
3D structure of target protein as predicted by I-TASSER server

**Figure 3:**
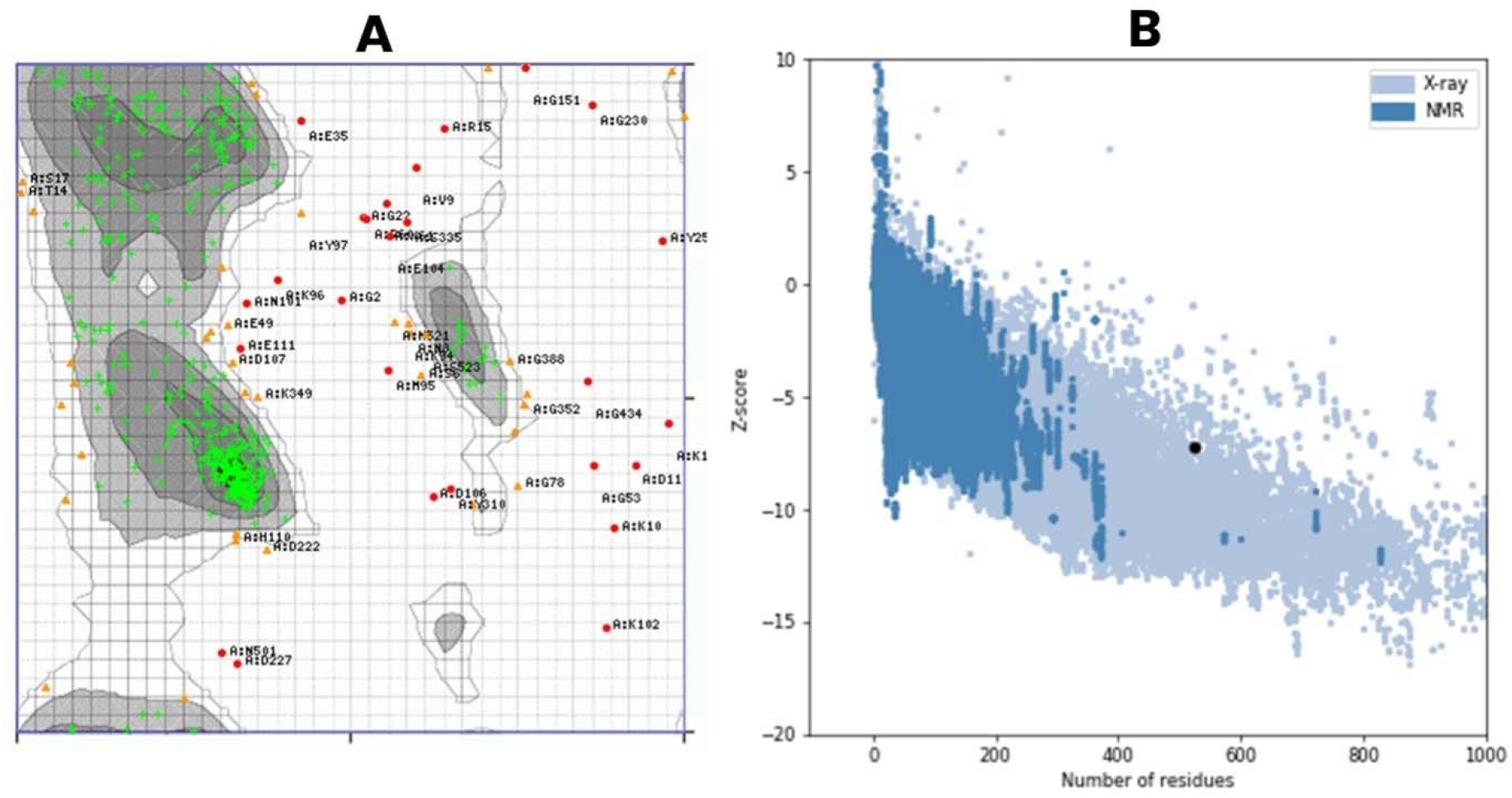
Validation of the tertiary structure of protein target. (A) Ramachandran plot statistics representing highly preferred region in green (87.93%), preferred region in Orange (7.0%) and questionable region in Red (4.98%). (B) ProSA web result representing Z-score of −7.17

**Figure 3:**
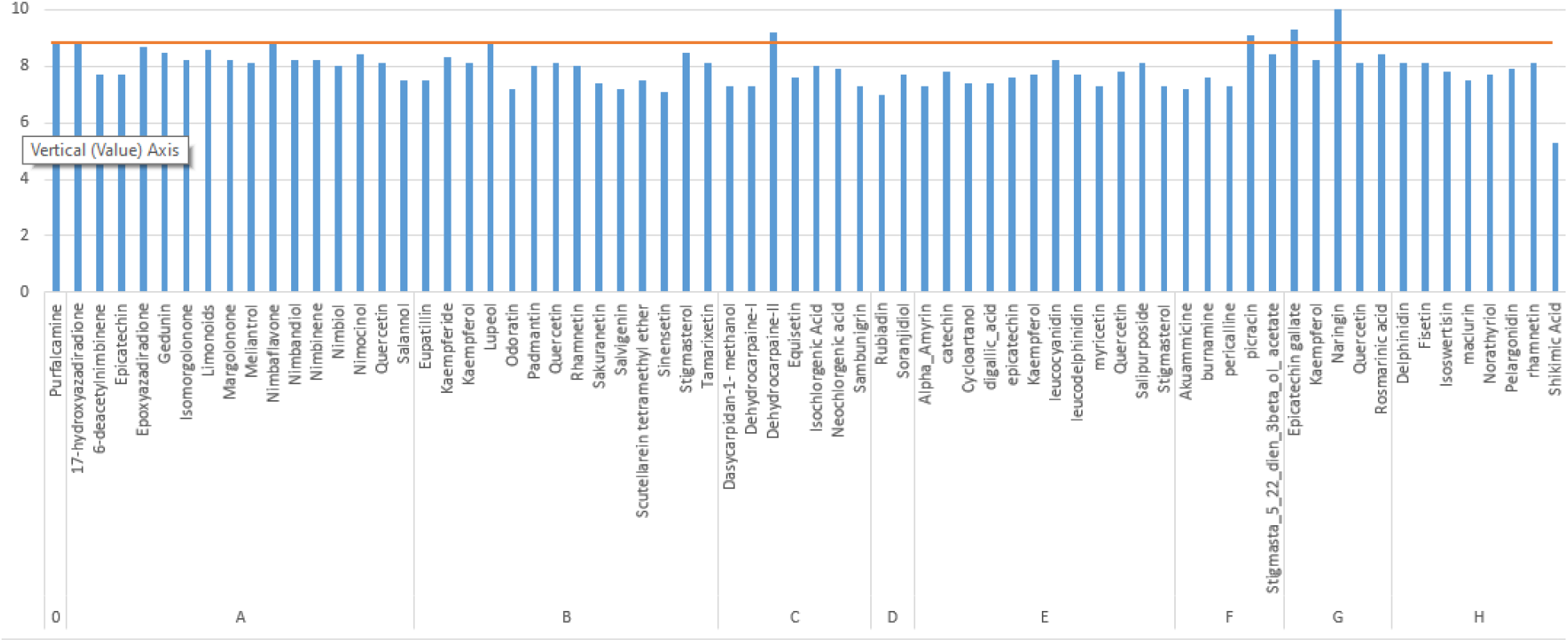
Graphical representation showing the binding affinities of selected phytochemicals and the reported PFCDPK1 inhibitor where group A represent *Azaradirachta indica, Group B* represent *Chromalaena Odorota*, Group C represent *Carica ppapaya*, Group D represent *Morinda lucida*, Group E represents *Anacardium ocidentale*, Group F represent *Pikralima nitida*, Group G represent *Citrus aurantofolia and* Group H represent *Magnifera indica*.

**Figure 4:**
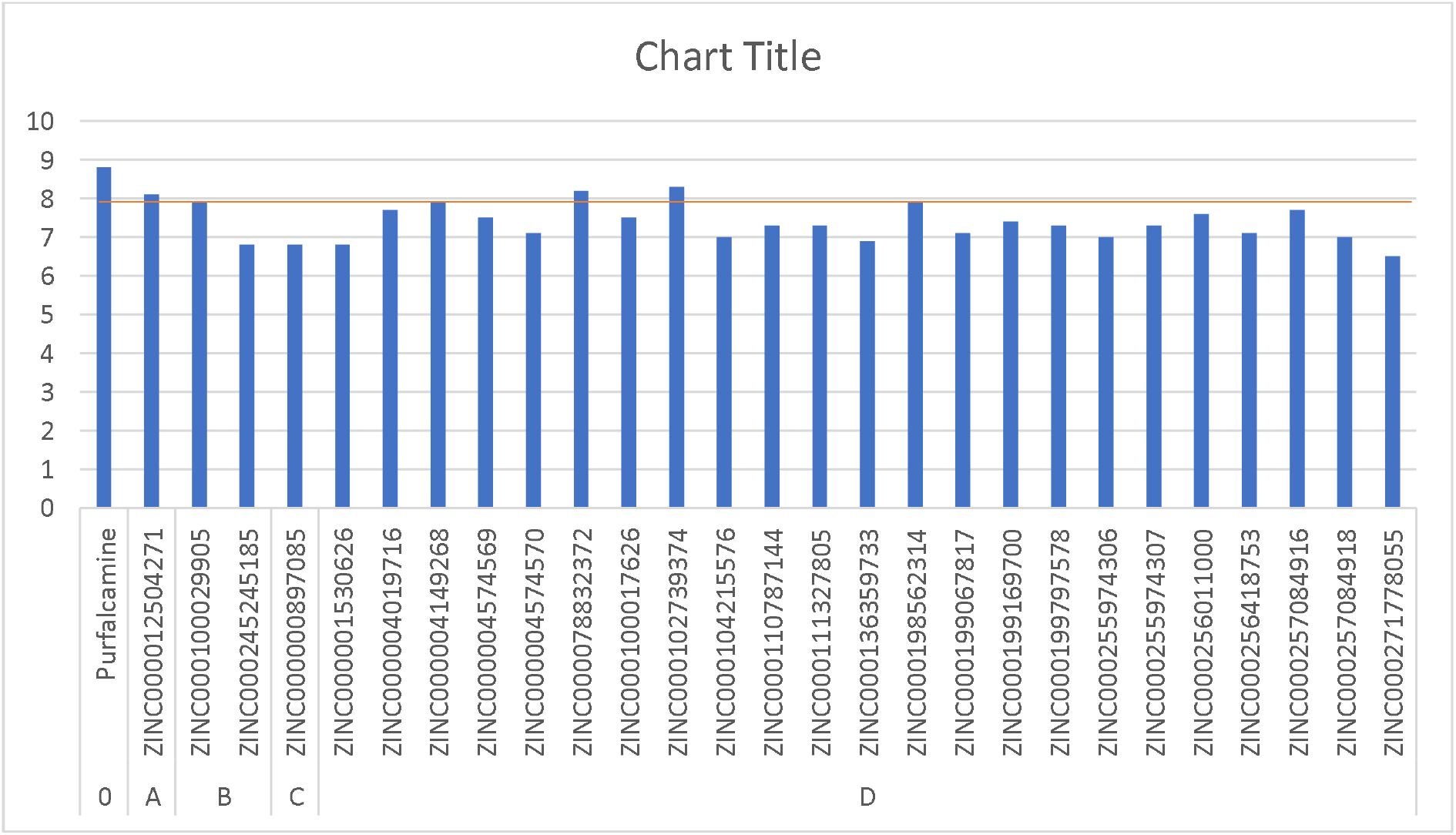
Graphical representation showing the binding affinities of selected pharmacophores and the reported PFCDPK1 inhibitor

### Determination of target protein active site

Analysis of the target protein 3D structure by CASTP showed that the protein’s pocket contains the following amino acids: HIS 125, PRO 126, ASN 127, GLU 146, TYR 148, LEU 153, GLN 156, ARG 160, PHE 163, CYS 166, ASP 167, ASN 170, ILE 171, LYS 173, GLN 174, SER 177, LEU 199, GLU 200, ASN 201, HIS 203, SER 204, LEU 203, LEU 206, ASN 207, ILE 208, LYS 209. The protein grid box dimensions were later determined using these amino acids.

### Ligand selection and preparation

PUBCHEM was used to retrieve a library of 472 ligands in total (39 ligands from *Chromalaena odorota*, 44 from *Azadirachta indica*, 120 from *Anacardium ocidentale*, 31 from *Citrus aurantofolia*, 68 from *Morinda lucida*, 13 from *Allium sativum*, 45 from *Carica papaya*, 45 from *Magnifera indica* and 67 from *Pikralima nitida*). Additionally, using ZINC15 server’s “for sale filter,” pharmacophoric modeling on commercial anti-malaria medications produced 385 pharmacophores (60 from Arthemeter, 8 from Atovarquine, 74 from Chloroquine, 95, Doxycycline, 1 from Lumenfantrine, 13 from Mefloquine, 25 from Primaquine, 107 from Quinine, and 2 from Tafenoquine). To find the ligands with the strongest hit and strong inhibitory capacity, all of the ligands were additionally analyzed using virtual screening. Supplementary Table 1 provides the ligand and pharmacophore CID.

### ADMET and drug-likeness profiling

The best ligands and pharmacophores with good ADMET (Adsorption, Distribution, Metabolism, Excretion, and Toxicity) result and drug-likeness properties were later selected after laborious assessment of the pharmacokinetic properties and drug-likeness of the retrieved ligands and pharmacophores (result available in supplementary table 2 and 3 respectively). In order to expedite the transformation of hits and lead compounds into approved candidates for therapeutic development, assessment of ADMET is a crucial component of the early stage of the drug discovery process. Additionally, *Pf*CDPK1 reported inhibitor (Purfalcamine) ADMET profiling was examined. The ADMET profiling shows that the reference compound (Purfalcamine) has a significant property of been hepatotoxic, hERG bloker, AMES toxic, carcinogenic, CYP3A4 inhibitor and with a very short clearance ability. Chosen compounds from antimalarial plants shows a promising ability of non-carcinogenic, mild-low ability of non-toxic, mild-low ability of non-hepatotoxic and a significantly low ability of been an hERG blocker. Nearly all the reported compounds have a good clearance ability, bioavailability and good intestinal absorption rate, Intrestingly, when critically accessed, the chosen compounds have a better therapeutical ability in terms of ADMET than the reference compound. Pharmacophores ADMET analysis showed a mild-low carcinogenicity property, very high intestinal absorption rate, high bioavailabilty, low CYP3A4 inhibitory power. Only two compounds show ability to inhibit p-glycoprotein while others shows a low property. Table 1 and 2 shows the ADMET result of selected compounds for both phytochemicals and pharmacophores respectively. The Lipinski rule of 5 law states compounds should not violate more than one of the following properties (1) Molecular weight is less or equal to 500, (2) LogP should be less or equal to 5, (3) Hydrogen bond donor should be less or equal to 5, (4) Hydrogen bond acceptor should not be more than 10 (Lipinski, 2004). The result obtained showed that only 4 compounds from both phytochemicals and pharmacophores violated more than one. Table 3 and 4 shows the drug-likeness properties (Lipinski Rule of five) of the selected phytochemicals and pharmacophores respectively.

**Table 1:**
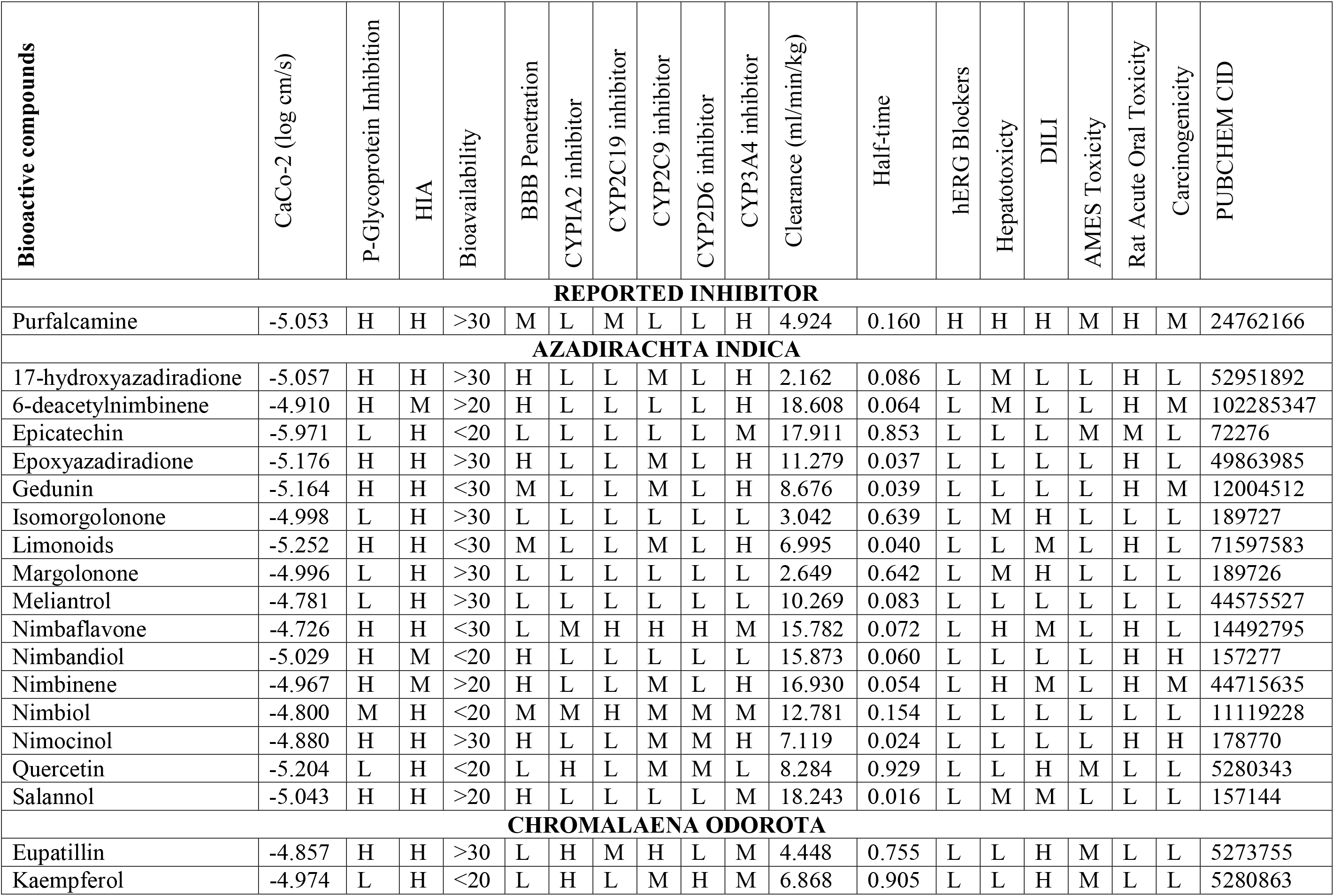
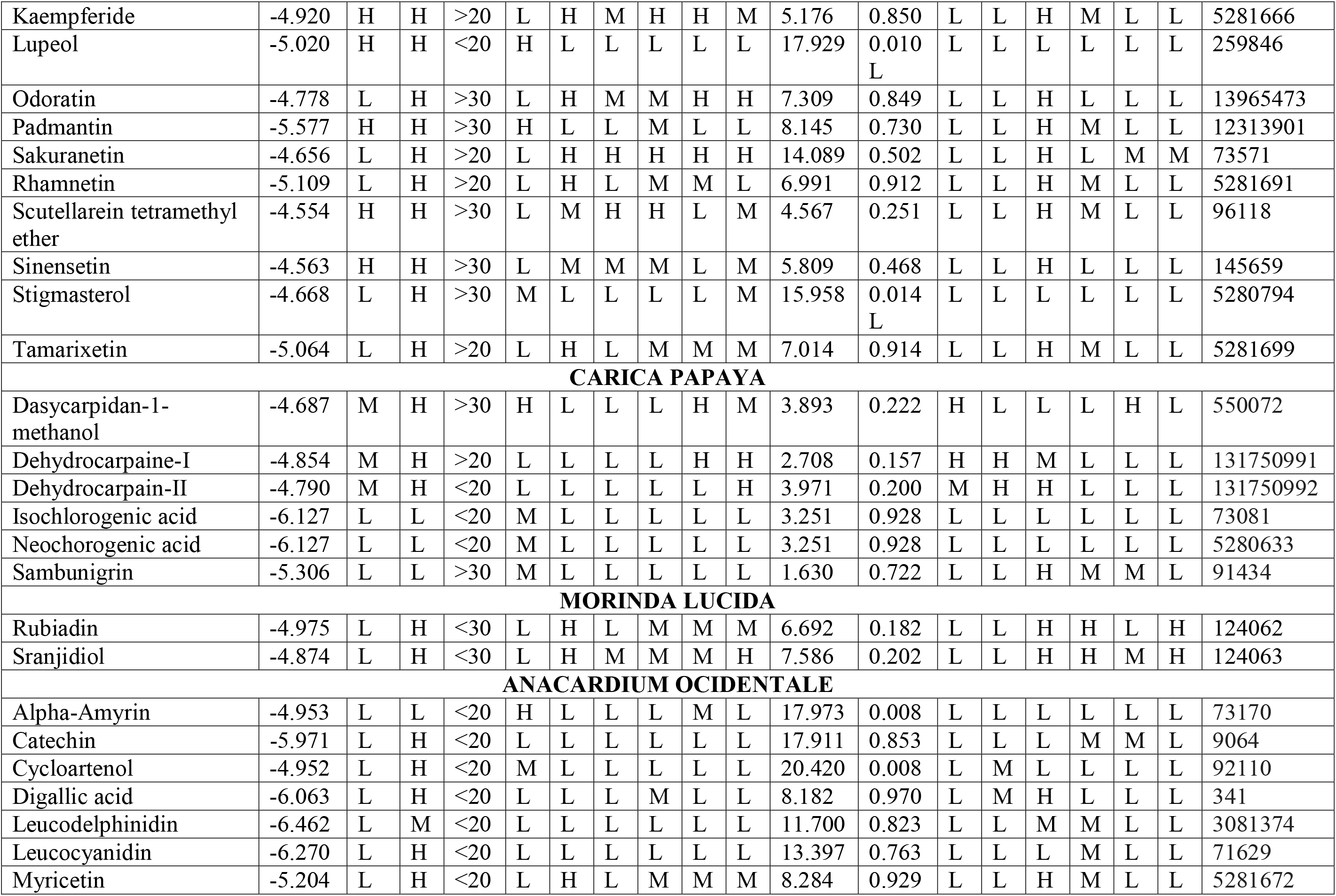
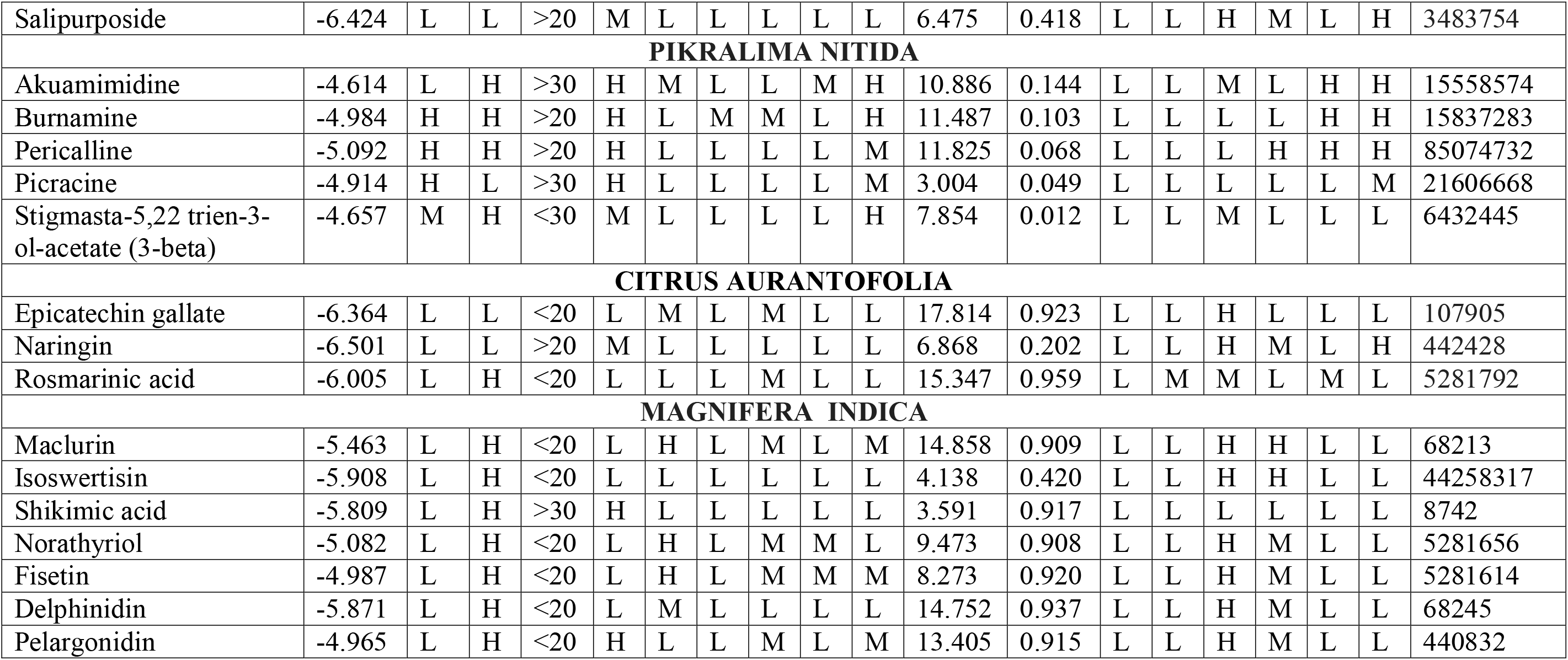
ADMET prediction of selected compounds per plant (Some compounds appeared in multiple plants analyzed, since ADMET property remains the same irrespective of the plant isolated from, they only appeared only once in our result).

**Table 2:**
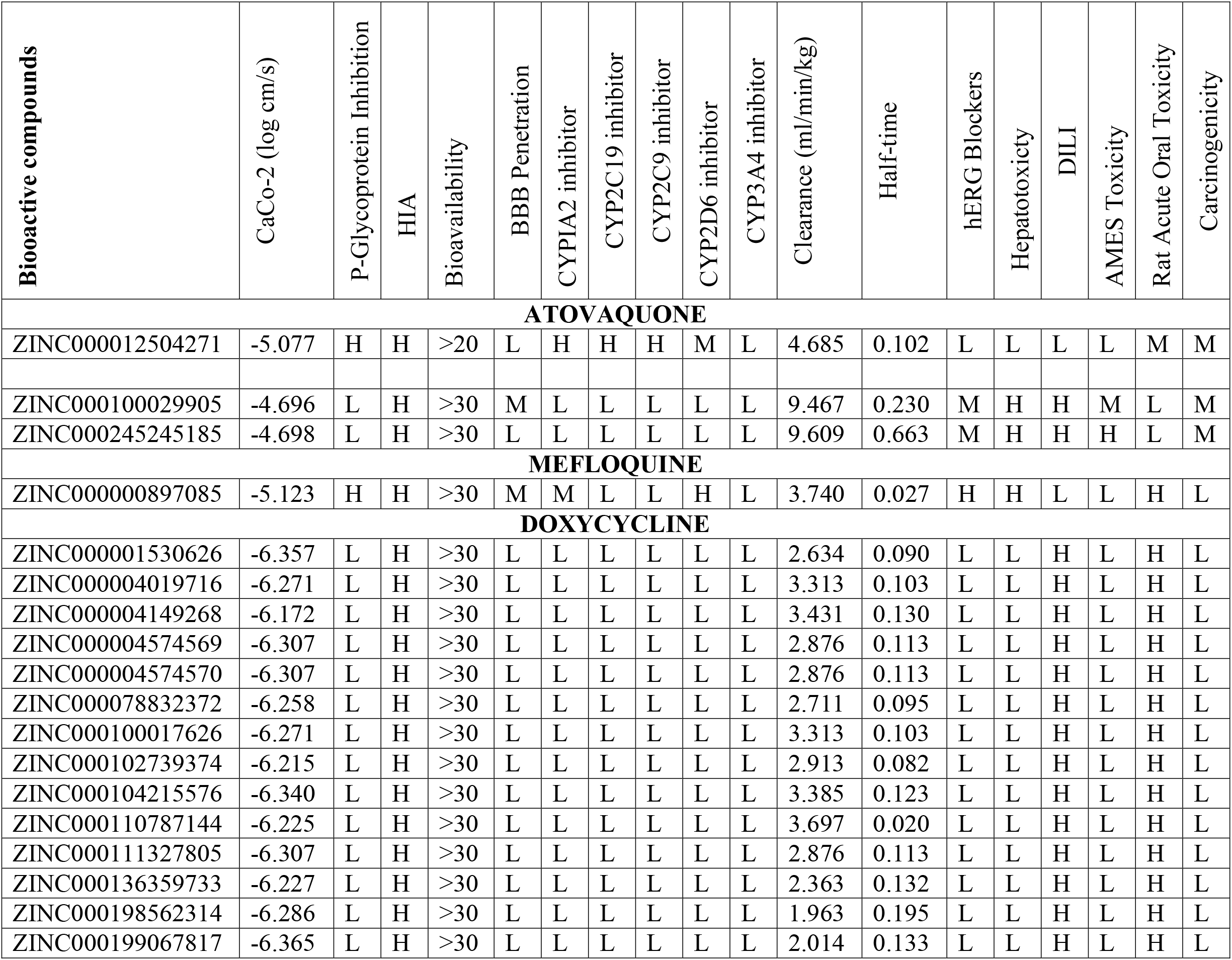
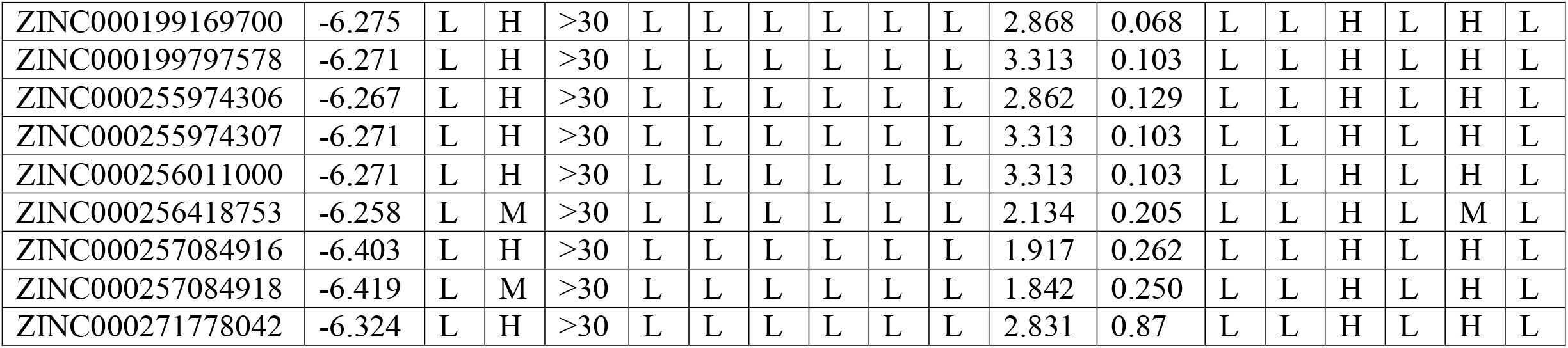
ADMET result for selected pharmaophores

**Table 3:**
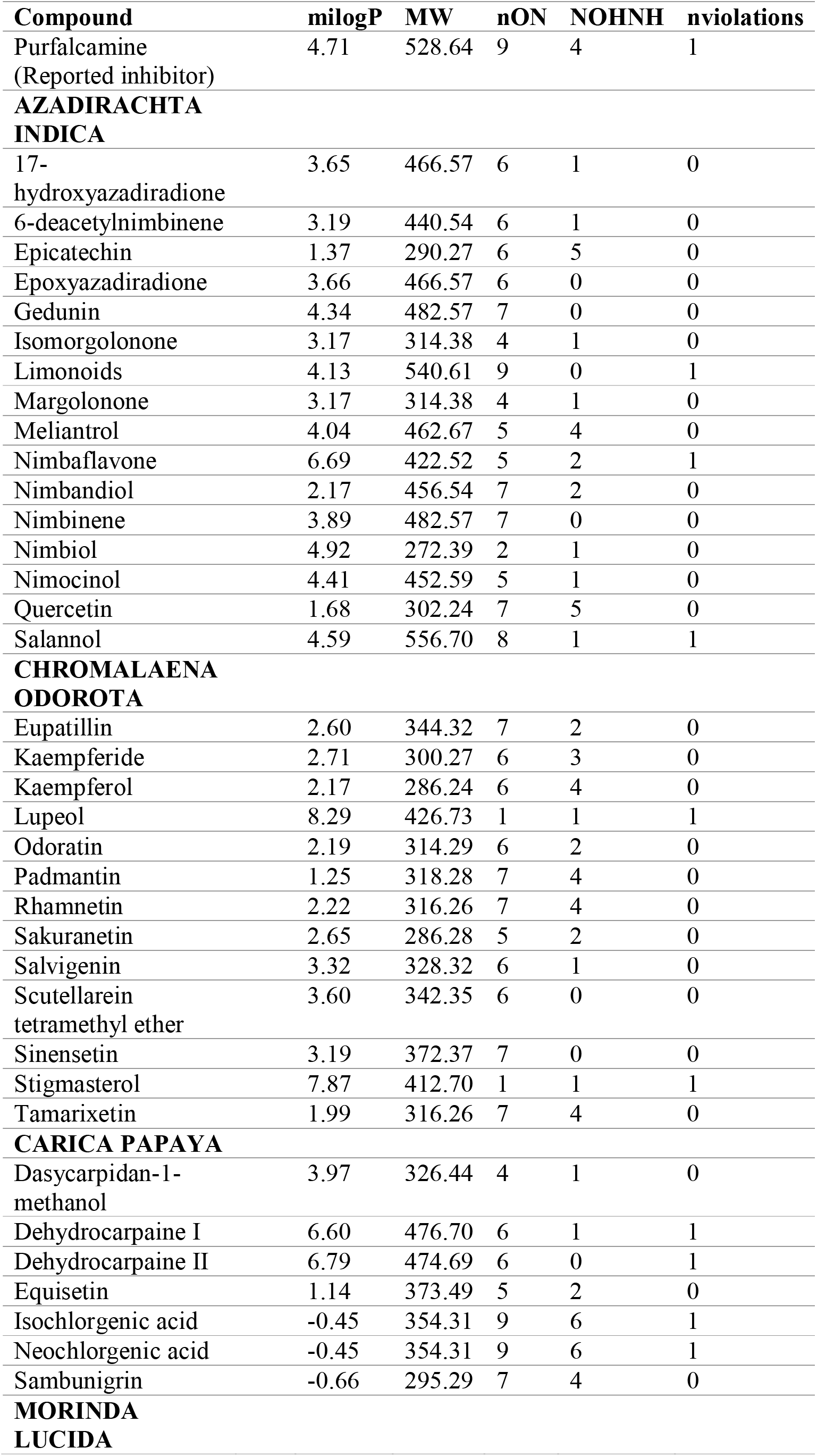
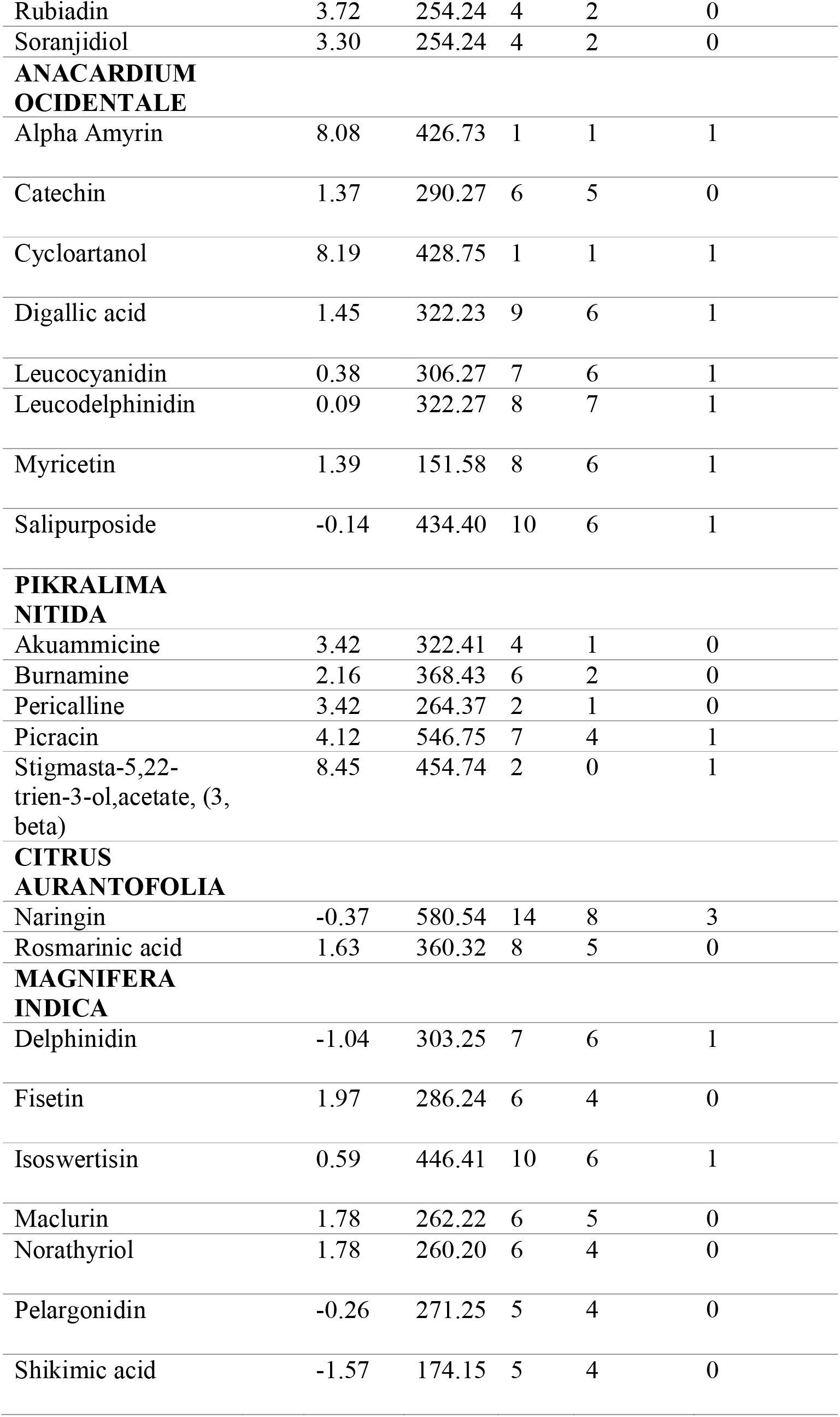
Drug-likeness (Rule of 5) parameters for selected phytochemicals

**Table 4:** Drug-likeness (Rule of 5) parameters for selected phytochemicals

| Compound | milogP | mw | nON | NOHNH | nviolations |
| --- | --- | --- | --- | --- | --- |
| **Atovaquine** |  |  |  |  |  |
| ZINC000012504271 | 4.96 | 366.84 | 3 | 1 | 0 |
| **Artemether** |  |  |  |  |  |
| ZINC000100029905 | 3.32 | 282.34 | 5 | 0 | 0 |
| ZINC000245245185 | 2.78 | 284.35 | 5 | 1 | 0 |
| **Mefloquine** |  |  |  |  |  |
| ZINC000000897085 | 4.24 | 378.32 | 3 | 2 | 0 |
| **Doxycycline** |  |  |  |  |  |
| ZINC000001530626 | -0.87 | 444.44 | 10 | 7 | 1 |
| ZINC000004019716 | -0.57 | 442.42 | 10 | 7 | 1 |
| ZINC000004149268 | 0.16 | 414.41 | 9 | 6 | 1 |
| ZINC000004574569 | -0.57 | 442.42 | 10 | 7 | 1 |
| ZINC000004574570 | -0.57 | 442.42 | 10 | 7 | 1 |
| ZINC000078832372 | 0.06 | 476.87 | 10 | 7 | 1 |
| ZINC000100017626 | -1.31 | 442.42 | 10 | 7 | 1 |
| ZINC000102739374 | 0.06 | 476.87 | 10 | 7 | 1 |
| ZINC000104215576 | -2.04 | 444.44 | 10 | 5 | 0 |
| ZINC000110787144 | 0.06 | 476.87 | 10 | 7 | 1 |
| ZINC000111327805 | -1.31 | 442.42 | 10 | 7 | 1 |
| ZINC000136359733 | -0.98 | 444.44 | 10 | 7 | 1 |
| ZINC000198562282 | -0.24 | 444.44 | 10 | 7 | 1 |
| ZINC000199067817 | -1.16 | 460.44 | 11 | 8 | 2 |
| ZINC000199169700 | 0.06 | 476.87 | 10 | 7 | 1 |
| ZINC000199797578 | -0.57 | 442.42 | 10 | 7 | 1 |
| ZINC000255974306 | -1.31 | 442.42 | 10 | 7 | 1 |
| ZINC000255974307 | -1.31 | 442.42 | 10 | 7 | 1 |
| ZINC000256011000 | -0.57 | 442.42 | 10 | 7 | 1 |
| ZINC000256418753 | -0.57 | 442.42 | 10 | 7 | 1 |
| ZINC000257084916 | -1.16 | 460.44 | 11 | 8 | 2 |
| ZINC000257084918 | -1.16 | 460.44 | 11 | 8 | 2 |
| ZINC000271778055 | -0.43 | 444.44 | 10 | 7 | 1 |

### Molecular Docking (Identification of best hit)

The retrieved substances were docked into the target protein’s active site, and the corresponding binding affinities of each compound are also listed in supplemental tables 4 and 5. Purfalcamine which was used as a standard compound showed −8.8KJ/mol binding affinity. All chosen compounds have binding affinity of higher or roughly near that of the standard. The final compounds chosen from ethnopharmaceutical investigations are listed in table 5 along with their amino acid interactions and binding affinities. Table 6 lists the pharmacophores that were selected. Graphs shown in figure 2 (Phytochemical) and 3 (Pharmacophore) shows the differences between selected compounds and Purfalcamine (a known *Pf*CDPK1 inhibitor) respectively. Their respective binding complex structures are provided in supplementary Figure 1.

**Table 5:** Binding affinity and molecular interaction of selected phytochemicals with PFCDPK1

| Compound | Binding Affinity (KJ/mol) | Hydrogen Bond | Other Bonds |
| --- | --- | --- | --- |
| **REPORTED INHIBITOR** |  |  |  |
| Purfalcamine | -8.8 | GLU 372, GLU 81 | LYS 375, GLU 200, LEU 371, LEU 206 |
| **AZADIRACHTA INDICA** |  |  |  |
| 17-hydroxyazadiradione | -8.8 | ASN 207, ASN 333 | LYS 375, LEU 371, ASN 170 |
| 6-deacetylnimbinene | -7.7 | ASN 207, ASP 379 | LYS 375, LEU 205, LEU 206, LYS 382, |
| Epicatechin | -7.7 | ASN 127, ASN 170, ASN 331, LYS 173 | LYS 282 |
| Epoxyazadiradione | -8.7 | LYS 382 | ASN 170, LYS 375, LEU 371, THR 378, ASP 379 |
| Gedunin | -8.5 | LYS 173 | LYS 382, LEU 206, LYS 375, LEU 205 |
| Isomorgolonone | -8.2 | ASN 201, LYS 382, GLU 372 | GLU 200, LEU 206 |
| Limonoids | -8.6 | ASN 170 | ASN 333, LYS 382, LEU 371, LYS 375 |
| Margolonone | -8.2 | ASP 287, TYR 258 | TYR 284, LYS 283, LYS 278, PHE 267, TYR 285 |
| Meliantrol | -8.1 | LYS 283 | TYR 284, PRO 266, TYR 264 |
| Nimbaflavone | -8.8 | ASN 333 | LYS 382, ASP 379, LYS 375, LEU 371, LEU 206, GLU 372 |
| Nimbandiol | -8.2 | ASN 170, GLU 200 | LYS 382, LEU 205, ASP 379, LEU 206 |
| Nimbinene | -8.2 | ASN 207, ASN 333 | LEU 206, LYS 382 |
| Nimbiol | -8.0 | LEU 377 | LYS 375, GLU 376, LYS 338 |
| Nimocinol | -8.4 | ASN 170, ASN 333 | LYS 382, LEU 206, LEU 371, LYS 375, LEU 205 |
| Quercetin | -8.1 | TYR 258, LYS 278, GLY 268 | LYS 283, TYR 284 |
| Salannol | -7.5 | ASN 170, ASN 333 | LYS 382, GLU 200, LYS 375, HIS 203 |
| **CHROMALAENA ODORATA** |  |  |  |
| Eupatillin | -7.5 | LYS 173, ASN 331 | ILE 332, LYS 383, LYS 382, ASP 385, GLY 388 |
| Kaempferide | -8.3 | TYR 258 | TYR 284, LYS 283, LYS 278 |
| Kaempferol | -8.1 | LYS 278, GLY 268 | LYS 283, TYR 284 |
| Lupeol | -8.8 | GLU 372 | LEU 205, LYS 382, LYS 375, LEU 206, LEU 371 |
| Odoratin | -7.2 | GLY 268, LYS 278 | LYS 283, TYR 284, TYR 264 |
| Padmantin | -8.0 | LYS 173, SER 177, LYS 383 | LYS 382, ASN 127 |
| Rhamnetin | -8.0 | LYS 173, ASP 385, ASN 207 | LYS 382 |
| Sakuranetin | -7.4 | LYS 283 | TYR 284, LYS 278, PRO 265 |
| Salvigenin | -7.2 | GLU 152, SER 64 | LEU 198, VAL 70, LYS 85, VAL 211, ASP 212 |
| Scutellarein tetramethyl ether | -7.5 | LYS 382 | LYS 173, LYS 383 |
| Sinensetin | -7.1 | ASN 170 | ASP 167, ARG 160, GLY 150, GLU 155 |
| Stigmasterol | -8.5 | GLU 372 | LYS 173, LYS 382 |
| Tamarixetin | -8.1 | PRO 265, GLY | LYS 278, LYS |
|  |  | 268 | 283, TYR 284 |
| **CARICA PAPAYA** |  |  |  |
| Dasycarpidan-1- methanol | -7.3 | TYR 285, PRO 266 | TYR 284, LYS 283, LYS 278 |
| Dehydrocarpaine-I | -7.3 | THR 378 | LEU 205, LYS 382, LYS 375 |
| Dehydrocarpaine-II | -9.2 | LYS 375 | - |
| Equisetin | -7.6 | ASN 207 | LYS 375, LEU 206, LYS 382, PRO 126 |
| Isochlorogenic Acid | -8.0 | ASN 127, LYS 173, GLY 388, ASN 331 | ASP 385, LYS 382 |
| Neochlorogenic acid | -7.9 | ASN 127, ASP 385, LYS 173, ASN 331 | LYS 382, LYS 383 |
| Sambunigrin | -7.3 | GLY 268, TYR 385, ASP 287 | TYR 284, LYS 278, LYS 283 |
| **MORINDA LUCIDA** |  |  |  |
| Rubiadin | -7.0 | ASN 170, ASN 331 | LYS 382, LYS 173 |
| Soranjidiol | -7.7 | TYR 328, GLY 390 | LYS 173, LYS 382, ASP 385, |
| **ANACARDIUM OCIDENTALE** |  |  |  |
| Alpha_Amyrin | -7.3 | GLU 372 | LYS 375, LEU 205, LEU 371, LEU 206, LYS 382 |
| catechin | -7.8 | ASN 127, SER 177, LYS 173 | LYS 386, LYS 382, ASP 385 |
| Cycloartanol | -7.4 | GLU 372 | LYS 375, LEU 206, LYS 382 |
| digallic_acid | -7.4 | SER 177, ASP 385, LYS 382, ASN 170, LEU 205, ASN 207 | - |
| epicatechin | -7.6 | LYS 173 | ASN 331, PRO 126, LYS 382 |
| Kaempferol | -7.7 | PRO 266, TYR 285 | PRO 265, TYR 264, GLY 268, LYS 278 |
| leucocyanidin | -8.2 | ASN 170, LYS 382, SER 177 | LYS 173 |
| leucodelphinidin | -7.7 | LYS 173, ASN 170, SER 177, ASP 385 | GLY 388 |
| myricetin | -7.3 | ASN 207, ASN 170, THR 378 | LYS 382, LEU 205 |
| Salipurposide | -8.1 | ASN 170, ASN 333, LEU 205, GLN 174, LYS 382 | LEU 371, GLU 372 |
| Stigmasterol | -7.3 | GLU 372 | LEU 206, LYS 382, LYS 173 |
| **PICRALIMA NITIDA** |  |  |  |
| Akuammicine | -7.2 | ARG 190 | ILE 99, ASP 227 |
| burnamine | -7.6 | ASN 207 | LEU 206 |
| pericalline | -7.3 | ASP 212 | LEU 198, LYS 85, ILE 129, VAL 211 |
| picracin | -9.1 | LYS 382, LEU 205 | GLY 388, LYS 173, LEU 371 |
| Stigmasta_5_22_dien_3beta_ol_acetate | -8.4 | - | LYS 375, LEU 371, LYS 382, LYS 173, SER 377 |
| **CITRUS AURANTOFOLIA** |  |  |  |
| Epicatechin gallate | -9.3 | ASN 170, ASN 333, LYS 173, GLY 390, SER 177, ASN 127 | LYS 382 |
| Naringin | -10.0 | ASP 385, LYS 173, ASN 170, ASN 207 | GLU 200, LEU 206, LEU 371, GLU 372 |
| Rosmarinic acid | -8.4 | GLY 388, ASN 127, ASN 207, ASN 333, GLU 200 | ASP 385, LYS 382, LYS 173 |
| **MAGNIFERA INDICA** |  |  |  |
| Delphinidin | -8.1 | GLY 388, LYS 382, ASN 170, LYS 173 | SER 177 |
| Fisetin | -8.1 | GLY 268, TYR 258, | LYS 283, LYS 278, PRO 266, TYR 284 |
| Isoswertisin | -7.8 | ASN 170, LEU 205, LYS 375, ASN 331 | LYS 173, LYS 382 |
| maclurin | -7.5 | ASP 385, LYS 173, LYS 386 | LYS 382 |
| Norathyriol | -7.7 | ASN 333, ASN 170, ASN 127, | LYS 173, LYS 382 |
|  |  | SER 177, ASP 385 |  |
| Pelargonidin | -7.9 | TYR 264, GLY 268 | LYS 278, TYHR 284, LYS 283 |
| Rhamnetin | -8.1 | LYS 173, ASP 385, SER 177, LYS 382 | - |
| Shikimic Acid | -5.3 | LYS 375, LEU 371 | LYS 382 |

**Table 6:** Binding affinity and molecular interaction of selected pharmacophores with PFCDPK1

| Compound | Binding Affinity | Hydrogen Bond | Other Bonds |
| --- | --- | --- | --- |
| **ATOVAQUONE** |  |  |  |
| ZINC000012504271 | -8.1 | LYS 173, ASN 333 | LYS 382, GLU 200, LEU 371 |
| **ARTHEMETER** |  |  |  |
| ZINC000100029905 | -7.9 | ASN 346 | ILE 450, MET 446, LEU 377 |
| ZINC000245245185 | -6.8 | ASN 333, ASP 379, THR 378 | LYS 382, LYS 375, LEU 371, PRO 126 |
| **MEFLOQUINE** |  |  |  |
| ZINC000000897085 | -6.8 | LYS 173, LYS 382, SER 177 | GLY 388, ASN 127, ASN 170, GLN 174, ASN 207 |
| **DOXYCYCLINE** |  |  |  |
| ZINC000001530626 | -6.8 | LYS 382, ASN 170, LYS 173 | ASN 207, LEU 205, ASN 333 |
| ZINC000004019716 | -7.7 | THR 378, LYS 375 | ASN 333, LEU 371, LEU 206, GLU 372, LYS 382, ASN 207, GLU 200 |
| ZINC000004149268 | -7.9 | LYS 173, ASN 333, TYR 328, ASN 170 | LYS 382 |
| ZINC000004574569 | -7.5 | LYS 382, ASN 333 | GLN 174, LEU 205, HIS 203, LEU 206, ASP 379 |
| ZINC000004574570 | -7.1 | TYR 328, ASN 170, LEU 205, LYS 382, ASN 207 | LEU 206, GLU 200, LEU 371 |
| ZINC000078832372 | -8.2 | AASN 331, LYS 173, TYR 328, ASN 333, ASP 385 | ALA 329, LYS 382, ASN 170, GLY 388 |
| ZINC000100017626 | -7.5 | ASP 287, PRO 266, GLY 268 | TYR 264, PRO 265, LYS 283, LYS 278, |
|  |  |  | PHE 267 |
| ZINC000102739374 | -8.3 | ASP 385, LYS 382, LYS 173 | ALA 329, LYS 326, GLY 388 |
| ZINC000104215576 | -7.0 | ASN 201, LYS 382, ASN 333, ASP 379, LYS 334 | LEU 206, LEU 371, THR 378, LYS 375 |
| ZINC000110787144 | -7.3 | ASP 385, ASN 387, LYS 386, LYS 173 | ALA 329, LYS 326, ASN 333 |
| ZINC000111327805 | -7.3 | LYS 382, ASN 333, THR 378, LYS 375 | GLU 200, LEU 371, LEU 206, ASP 379 |
| ZINC000136359733 | -6.9 | THR 339, ASN 346, ASP 447 | MET 446 |
| ZINC000198562314 | -7.9 | ASN 170, ASN 333 | LYS 382, LYS 375 |
| ZINC000199067817 | -7.1 | LYS 85, GLU 152, ASP 212 | - |
| ZINC000199169700 | -7.4 | ASP 385, LYS 382, ASN 333 | LYS 173, ILE 325, LYS 326, GLY 388, ILE 332 |
| ZINC000199797578 | -7.3 | ASN 333, LYS 382 | LEU 206, LEU 371 |
| ZINC000255974306 | -7.0 | LYS 375, THR 378, LYS 382, GLU 200 | LEU 206, LEU 205 |
| ZINC000255974307 | -7.3 | ASN 333 | THR 378, ASN 207, LEU 205, LEU 206, GLU 200, LEU 371 |
| ZINC000256011000 | -7.6 | GLY 268 | LYS 278, PHE 267, LYS 283 |
| ZINC000256418753 | -7.1 | LYS 382, ASN 333, LYS 173, ASN 331 | LYS 326 |
| ZINC000257084916 | -7.7 | LYS 283, TYR 284 | TYR 284 |
| ZINC000257084918 | -7.0 | ASP 447, ASN 346 | LYS 338, MET 446 |
| ZINC000271778055 | -6.5 | LYS 382, ASN 207, LEU 205, ASN 333 | GLU 200, LEU 371, ASN 170 |

## DISCUSSION

*Plasmodium falciparum* is a major causative agent of pernicious human malaria, which infects the mosquito and the human host. Hepatocytes and erythrocytes are the two types of human host cells that *Plasmodium* infects. It also encounters other host conditions, which it successfully survives to spread (Kumar et al., 2021). Sadly, signs of antimalarial medication resistance are putting continuing control efforts at risk (Flaherty et al., 2019; Large et al., 2013). Future control efforts will depend on the creation of novel antimalarial medications as well as the identification and characterization of novel Plasmodium protein targets (Bansal et al., 2017; Flaherty et al., 2019; Iyer et al., 2018; Large et al., 2013). Right here, we target *Pf*CDPK1 that has been found in sporozoites, sexual stages, and asexual blood stages. The gene appears to be expressed in *Plasmodium falciparum* asexual blood stages from the start of schizogony (at around 30 h post invasion), and the protein builds up in the cell until the end of the cycle (Green et al., 2008; Holder et al., 2012; Large et al., 2013). According to research by Bansal et al (2013), this protein is able to phosphorylate parts of the molecular motor that propels parasite penetration of red blood cells and is implicated in parasite motility and host cell invasion.

A biological target is digitally docked with collections of millions of chemicals as part of the *in silico* technique known as virtual screening, and the tightness of the fit is then assessed (scoring). Virtual screening minimizes the number of compounds that must be tested in order to find early hits and provides speedy evaluation of large libraries (Kontoyianni, 2017; Pinzi & Rastelli, 2019; Yuriev & Ramsland, 2013). The creation of new, promising chemicals has greatly benefited from the merging of computational and experimental methodologies. Molecular docking techniques, which are widely utilized in contemporary drug design, examine the ligand conformations taken within the binding sites of macromolecular targets (Ferreira et al., 2015).

In the current investigation, we used molecular docking, ADMET analysis, and drug-likeness profiling to virtually screen 856 small compounds for potential PfCDPK1 inhibitors. The target molecule’s structural integrity determines how effective the outcome will be. The Ramachandran plot server and ProSA web were used to validate the modeled enzyme target. Analysis of the results showed that the modeled structure is trustworthy and may be utilized for virtual screening (Amino acid in very favorable region equals 87.13% and z-score of −7.17). There have been far too few reports of *Pf*CDPK1 inhibitors (Lemercier et al., 2009). Here, we explore these small compounds more thoroughly and with greater therapeutic effectiveness and drug-likeness. Purfalcamine (a putative inhibitor of *Pf*CDPK1) and *Pf*CDPK1 were found to have a virtual hit, with the analysis of the result showing −8.8Kcal/mol, two amino acids participating in the hydrogen bond interaction and four other amino acids participating in other bonds. This interaction shows that the target protein and the ligand have a high binding affinity.

According to ADMETlab, a clearance value > 15 ml/mm/kg is regarded high, 5-14.9 ml/mm/kg is considered moderate, and <5 ml/mm/kg is considered low. A good therapeutic medication demonstrated ease of excretion to prevent toxin buildup (Garza et al., 2022). The purfalcamine clearance score was 4.924 ml/mm/kg, indicating a limited clearance capacity that can invariably lead to toxin accumulation and, as a result, to hepatotoxicity and drug-induced liver injury (DILI). Due to its propensity to inhibit CYP3A4, it has a limited clearance capacity. Over 50% of drug metabolism is carried out by the cytochrome P450 family enzymes, which belong to the CYP3A4 class (Garza et al., 2022). In addition to having a significant AMES toxicity effect, purfalcamine also has the potential to cause cancer. Purfalcamine’s drug-likeness assessment revealed one significant violation (Molecular weight > 500 kDa). The Lipinski rule of five states that a compound’s molecular weight must be less than 500kDa. A decrease in absorption is caused by an increase in molecular weight because the concentration of the molecule at the intestinal epithelium’s surface decreases (Tripathy, 2019).

*Alium sativum* compounds were eliminated from further consideration because they had a very low binding affinity to the target enzyme’s active site when compared to the other therapeutic plant phytochemicals. Compounds from *Azadirachta indica* were discovered to interact strongly with the target protein. It was discovered that asparagine (ASN) molecules contributed roughly 68% of the hydrogen bonds to the chosen compounds. Typically, hydrogen bonds are referred to as “enhancers of protein-ligand binding” (Salentin et al., 2014). Therefore, it might imply that their high binding affinities were influenced by the robust hydrogen bonds. Additionally, we noticed that around 87% of the compounds we chose have the lysine (LYS) residue conserved in other bonds. We provided evidence to support our claim that this residue may be significantly influencing the reported complexes’ high binding affinities. Also noted was the presence of LYS375, which is also seen in other bonds, in the interaction between the target protein and purfalcamine. Compared to purfalcamine, all of the compounds that were chosen from this plant have improved ADMET profiles, but only two of them—17-hydroxyazadiradione and nimbaflavone—are able to bind with an affinity of −8.8 kJ/mol.

More LYS residue was also seen in the phytochemicals from *Chromalaena odorata*, *Carica papaya*, *Morinda lucida*, *Anacardium ocidentale*, *Picralima nitida*, *Citrus aurantofolia*, and *Magnifera indica*. This suggests that LYS residue is always significant and helps explain why these phytochemicals have a higher binding affinity. We observed that Dehydrocarpaine-II (from *Carica papaya*), Picracin (from *Picralima nitida*), Epicatechin gallate and Naringin (from *Citrus aurantofolia*) has higher binding affinity for the active pocket of *Pf*CDPK1 (−9.2, −9.1, −9.3, −10.0 Kcal/mol) compared to purfalcamine. These chemicals can be a better inhibitory effector to the protein target. Comparing the ADMET profiles of all the phytochemicals used in this investigation to the *Pf*CDPK1 inhibitory benchmark, they are all better.

Pharmacophoric modeling was carried out on nine (9) reported anti-malarial medicines in order to further explore for substances that have good potential inhibitory action on the target enzyme. Only four (4) of these groups - Atovaquone, Arthemether, Mefloquine, and Doxycycline - were discovered to have a higher affinity for the target protein. Pharmacophores for the antibiotic doxycycline were discovered to exhibit high binding affinity for *Pf*CDPK1. The fact that these pharmacophores have a relatively low molecular weight (500kda) might suggest their easily digestible character. Similarly, the pharmacophoric compounds showed low level of been carcinogenic, good bioavailability and intestinal absorption rate. This implies they have the ability to reach their target site before they become degraded. Additionally, none of the pharmacophores used in this investigation had any capacity to inhibit CYP314 inhibitor, which reduces their potential to be hepatotoxic. However, our result points to 6 compounds from the pharmacophores with ZINC ID-ZINC000012504271, ZINCC000100029905, ZINC000004149268, ZINC000078832372, ZINC000102739374 and ZINC000198562314. These compounds exhibited better binding affinities of −8.1, −7.9, −7.9, −8.2, −8.3, −7.9 KJ/mol respectively.

## CONCLUSION

With minimal pharmacokinetic and pharmacodynamic adverse effects, this work aims to find more potential small molecule ligands with better *Pf*CDPK1 inhibitory effects. Using various virtual screening techniques, 856 compounds in total were tested. In the end, 88 compounds were chosen because they were more effective than purfalcamine. We hereby propose that these compounds may be powerful enough to inhibit *Pf*CDPK1. Interestingly; our findings emphasize 17-hydroxyazadiradione, Nimbaflavone, Dehydrocarpaine-II, Picracin, Epicatechin gallate, Naringin, ZINC000012504271, ZINCC000100029905, ZINC000004149268, ZINC000078832372, ZINC000102739374 and ZINC000198562314 Further complex computational modeling and *in vivo* investigations is now necessary to evaluate these compounds and determine whether the chosen compounds are suitable therapeutic candidates.

## AKNOWLEDGEMENT

We acknowledge and appreciate all members of Computational Molecular Biology and Drug Discovery Laboratory, Department of Biochemistry, Ladoke Akintola University of Technology. Also, we appreciate the management and staffs of Helix Biogen Institute for providing technical support throughout the process of this research.

## AUTHOR’S CONTRIBUTION

TIA-Project conceptualization and design, TOO, BOQ-Data retrieval, analysis and wrote first draft manuscript, PMO-Data retrieval and analysis, EKO-Manuscript editing and review. All authors participated in the review of the final edition of the manuscript.

## DECLARATION OF COMPETING INTEREST

The authors have declared no known competing financial interest or personal relationship that might have appeared to influence the work reported here.

## FUNDING DECLARATION

We declare that we receive no funding throughout the research of this work.

## Notes

### Competing Interest Statement

The authors have declared no competing interest.

